# An immuno-suppressive aphid saliva protein is delivered into the cytosol of plant mesophyll cells during feeding

**DOI:** 10.1101/071811

**Authors:** Sam T. Mugford, Elaine Barclay, Claire Drurey, Kim C. Findlay, Saskia A. Hogenhout

## Abstract

Herbivore selection of plant hosts and plant responses to insect colonization have been subjects of intense investigations. A growing body of evidence suggests that for successful colonization to occur, (effector/virulence) proteins in insect saliva must modulate plant defense responses to the benefit of the insect. A range of insect saliva proteins that modulate plant defense responses have been identified, but there is no direct evidence that these proteins are delivered into specific plant tissues and enter plant cells. Aphids and other sap-sucking insects of the order Hemiptera use their specialized mouthparts (stylets) to probe plant mesophyll cells, until they reach the phloem cells for long-term feeding. Here we show by immunogold-labeling of ultrathin sections of aphid feeding sites that an immuno-suppressive aphid effector localizes in the cytoplasm of mesophyll cells near aphid stylets, but not in cells further away from aphid feeding sites. In contrast, another aphid effector protein localizes in the sheaths composed of gelling saliva that surround the aphid stylets. Thus, insects deliver effectors directly into plant tissue. Moreover, different aphid effectors locate extracellularly in the sheath saliva or are introduced into the cytoplasm of plant cells.

## Introduction

Insect herbivores likely modulate a range of plant processes in order to successfully colonize their host plants. A growing body of evidence suggests that the saliva of herbivorous insects contains virulence factors (effectors) that interfere with host plant defences and facilitate colonisation (Musser et al. 2002; Will et al. 2007; Mutti et al 2008; Bos et al. 2010; Stuart et al. 2012; Atamian et al. 2013; Pitino and Hogenhout 2013; Chaudhary et al., 2014; Elzinga et al. 2014; Guo et al. 2014; Rodriguez et al. 2014; Naessens et al. 2015; Wang et al. 2015) Sap-sucking insects of the order Hemiptera, such as aphids, whiteflies, leafhoppers, psyllids and planthoppers, are stealthy feeders. These insects possess piercing-sucking mouthparts, consisting of a pair of stylets, which navigate to vascular elements for long-term feeding. Hemipterans cause direct feeding damage to plants and also transmit the majority of plant viruses and several plant pathogenic bacteria (Hogenhout et al 2008; Guerrieri and Digilio 2008; Orlovskis et al. 2015). The green peach aphid (GPA) *Myzus persicae* alone transmits over a 100 plant viruses.

The feeding behaviour of aphids has been studied extensively. While navigating to the phloem, in the pathway phase, aphid stylets probe various plant cells of the mesophyll (Tjallingii 2006). During a probe the aphid stylets penetrate cells, deposit watery saliva inside, and acquire cell contents (Martín et al. 1997). The pathway phase continues until the aphid reaches a phloem sieve-element cell, where they remain to feed often for many hours. The phloem feeding phase starts with release of saliva followed by extended periods of phloem sap acquisition that are occasionally interrupted with short salivation periods (Tjallingii and Esch 1993). Whereas it has been shown that the probing of mesophyll cellsis responsible for the transmission of a diversity of viruses by aphids, the biological significance of this behaviour to the aphids themselves is not fully understood.

Aphids produce different types of saliva: ‘gelling’ saliva that forms a sheath around the stylets that is thought to have a protective function (Miles 1999), and ‘watery’ saliva that is thought to be injected into plant cells during the pathway and phloem feeding phases (Tjallingii and Esch 1993; Martín et al. 1997). A number of proteins are detected in aphid saliva (Ramsey et al. 2007; Harmel et al. 2008; Carolan et al. 2009; Cooper et al. 2010; Nicholson et al. 2012, 2014; Rao et al. 2013; Vandermoten et al. 2014; Chaudhary et al. 2015). Aphid saliva proteins have also been detected in lysate of plant tissues previously exposed to aphids (Mutti et al. 2008; Naessens et al. 2015). Some components of aphid saliva trigger plant defence responses that are characteristic of pattern-triggered immunity (PTI), including reactive oxygen species (ROS) bursts and callose deposition, that require the plant membrane-associated receptor-like kinase (RLK) BRASSINOSTEROID INSENSITIVE 1-ASSOCIATED KINASE (BAK1) (Chaudhary et al. 2014; Prince et al. 2014). BAK1 is a key regulator of several cell membrane-localized pattern recognition receptors (PRRs), which mediate the first step of the plant defence response via the recognition of pathogen-associated molecular patterns, such as flg22 (Chinchilla et al. 2007; Chaudhary et al. 2014).

Aphid saliva contains effectors that suppress plant defence responses (Will et al. 2007; Mutti et al 2008; Bos et al. 2010; Atamian et al. 2013; Pitino and Hogenhout 2013; Chaudhary et al., 2014; Elzinga et al. 2014; Guo et al. 2014; Rodriguez et al. 2014; Naessens et al. 2015; Wang et al. 2015). Mp10 is one of a number of candidate effector proteins that have been identified and was found to suppress the plant ROS burst in response to flg22 (Bos et al. 2010; Rodriguez et al. 2014). Mp10 and another candidate effector protein expressed in GPA salivary glands, MpOS-D1, belong to the chemosensory protein (CSP) family (Bos et al. 2010; Zhou et al. 2010). Other aphid effector candidates, such as MpPIntO1 and MpC002, promote aphid colonization (Mutti et al 2008; Pitino and Hogenhout 2013; Pitino et al. 2011; Coleman et al. 2015) and are found in aphid saliva and/or in extracts of aphid-exposed leaves (Mutti et al 2008; Harmel et al. 2008). However, evidence that insects deliver effectors directly into plant cells is currently lacking.

How pathogens and pests deliver effectors to the appropriate site of action has been a major research topic of many research groups (Dodds and Rathjen 2010). Gram-negative bacterial pathogens deliver their effectors via specialized mechanisms, such as type III secretion systems (Gopalan et al. 1998). For fungi and oomycetes it was discovered that effectors require conserved sequences, such as an RXLR motif, at their N-termini for entering the plant host, but the mechanisms underlying their delivery is not fully understood (Petre and Kamoun 2014). It is as yet unclear how, where and when aphids deposit their effectors in the plant; they may do so via the ‘gelling’ saliva to embed the proteins within the sheaths surrounding the stylets in the apoplastic space of plant tissue, or via ‘watery’ saliva to introduce them into plant cells. In addition, aphids may secrete different effectors during the pathway and phloem feeding phases.

Here we studied ultra-thin sections of plant tissues containing tracks of aphid stylets and used antisera raised to the aphid effectors Mp10, MpOS-D1, MpC002 and MpPIntO1 for immuno-gold labelling of the sections. Antisera of Mp10, but not of MpOS-D1, labelled the cytosol and chloroplasts of plant mesophyll cells adjacent to aphid stylet tracks, whereas antisera to MpPIntO1 and MpC002 labelled the aphid stylet sheaths. These data indicate that aphid effectors are delivered into the cytosol of plant cells during probing in the pathway phase, whereas other effectors are embedded within the sheaths that surround the stylets in the apoplastic space of mesophyll tissue.

## Results

### Candidate effector proteins are detected in total protein extracts of aphid-exposed leaves

Affinity purified antisera were raised against recombinant GPA candidate effectors and examined for specificity and background levels in aphid and plant extracts by protein immuno-blotting. The antisera to Mp10, MpOSD1, MpPIntO1 and MpC002 detected bands of the predicted sizes in total protein extracted from whole aphids (Fig. 1). Antisera to the CSPs Mp10 and MpOS-D1 detected Mp10 and MpOS-D1, respectively, down to the ng level (Supplementary Fig. S1), indicating that the antisera to these proteins are specific and sensitive. The antisera to all four effectors detected proteins of the predicted sizes in extracts of *A. thaliana* leaves previously exposed to GPA (Fig. 1), but not in unexposed leaves. These bands were not detected in blots probed with corresponding pre-immune sera (Supplementary Fig. S2). The Mp10 and MpOS-D1 antibodies and pre-immune sera reacted with some non-specific bands of different sizes in the aphid-infested and uninfested plant tissues, but the bands corresponding to the Mp10 and MpOS-D1 proteins were only detected in infested tissues and not with the pre-immune sera (Fig. 1, Supplementary Fig. S2). Thus, GPA delivers all four effectors into plant tissues during feeding.

**Figure 1.**
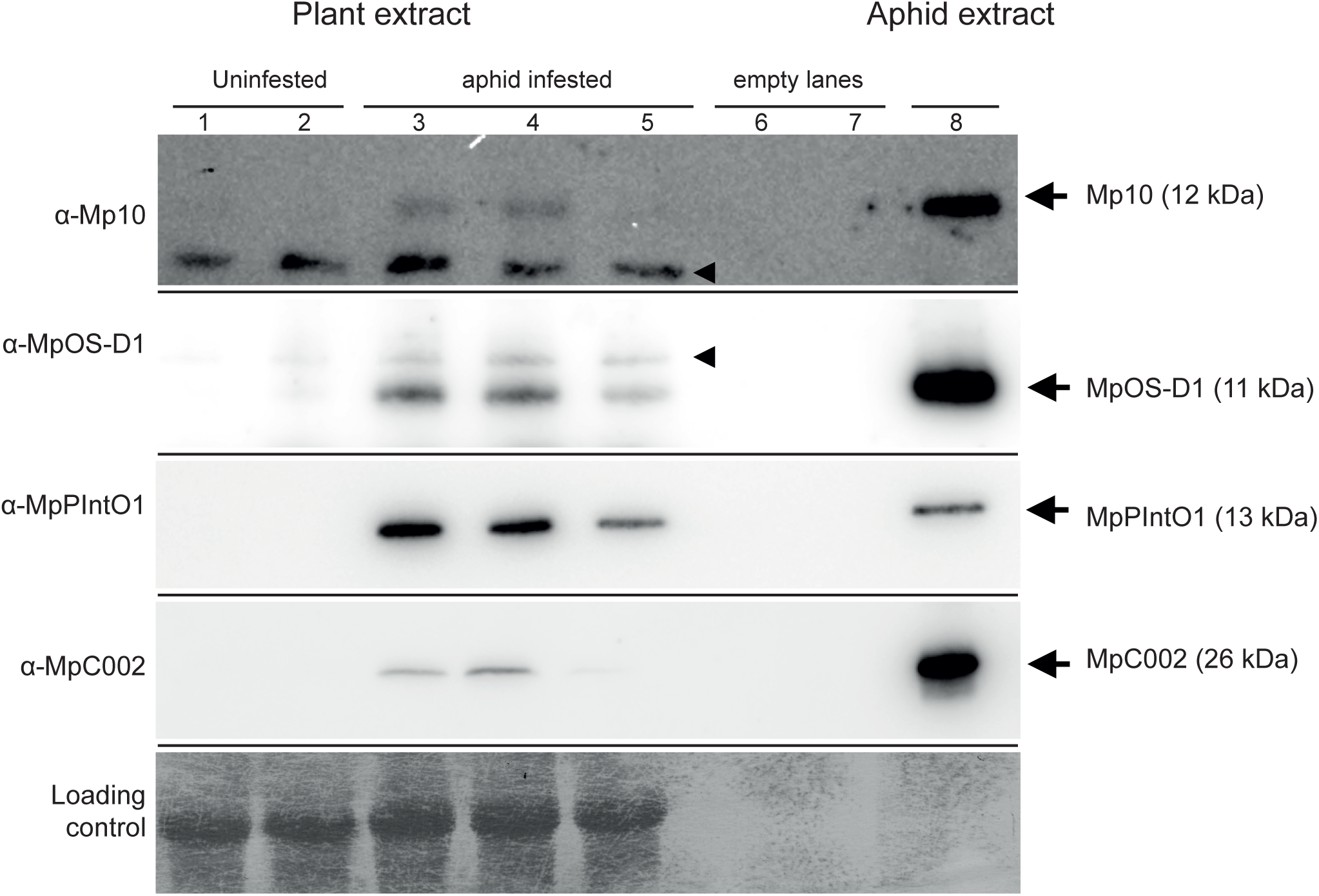
Aphid salivary proteins are detected in extracts of aphid-infested host plant tissue. Total protein extracts from whole aphids (GPA) (lane 8) and *A. thaliana* rosettes uninfested (lanes 1-2, two different plants) or infested with GPA (lanes 3-5, three different plants) were separated by SDS-PAGE, transferred to nitrocellulose membranes and probed with antisera raised against the aphid proteins Mp10, MpOS-D1, MpPIntO1 and MpC002 as shown at the left of the blots. The bottom panel shows an amido-black stained loading control. Arrows at right point to bands of predicted sizes for the effectors, as indicated. Arrowheads indicate non-specific bands detected by antibodies.

### The aphid effector protein Mp10 localizes to the cytoplasm and chloroplasts of mesophyll cells adjacent to aphid feeding sites

To investigate if GPA delivers effectors in or near feeding sites, semi-thin sections of GPA-infested leaf samples were stained with toluidine blue to localize GPA stylet tracks (Fig. 2A). Ultra-thin sections of the same sample were then used for immunogold-labelling (IGL) with the Mp10 antiserum (Fig. 2B-D). Controls included the GPA-infested *A. thaliana* leaf samples sections incubated with pre-immune sera and sections of uninfested *A. thaliana* leaf samples incubated with antisera (Fig, 3, 4 and Supplementary Fig. S2).

**Figure 2.**
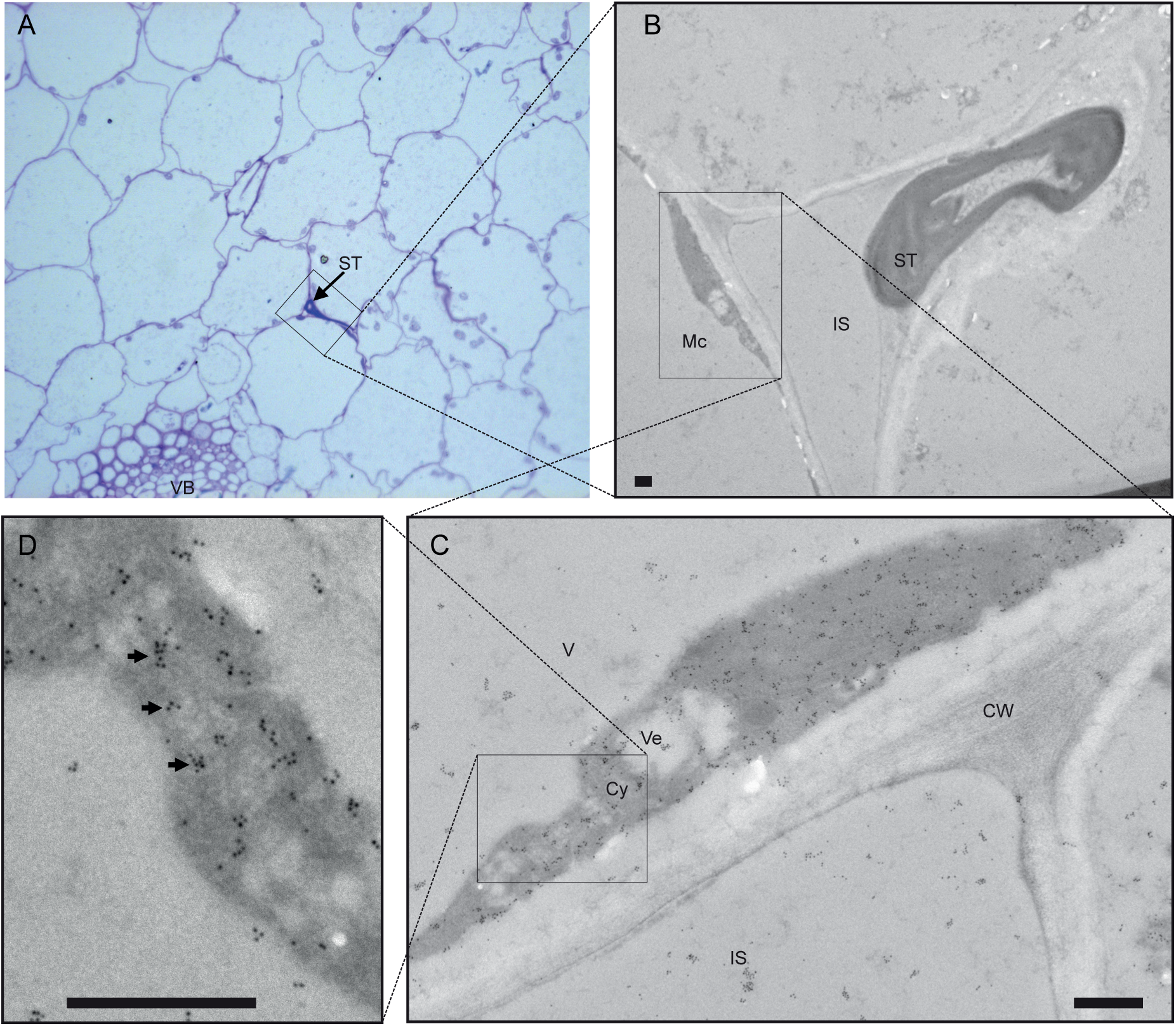
Mp10 protein is detected in the cytoplasm of mesophyll cells near an aphid stylets track. (A): Semi-thin section (0.5 µm) from an aphid-infested *A. thaliana* leaf stained with toluidine blue. Aphid stylets tracks (ST) are visible between mesophyll cells (Mc) and a vascular bundle (VB). (B-D): Immuno-gold labeling of ultra-thin sections (90 nm) of the tissue shown in (A) with anti-Mp10 (1:10 dilution) revealed a high density of gold particles (arrows in D) in the cytoplasm (Cy) of mesophyll cells (Mc). No obvious labeling was observed of aphid stylets (St), cell wall (CW), intercellular space (IS), mesophyll cell vacuole (V) and inside vesicles (Ve). Images shown in B to D are sequentially higher magnifications of the same area of a cell adjacent to the stylets track. Scale bars, 500 nm.

IGL with Mp10 antiserum (dilutions 1:10 and 1:100) showed high labelling density in cytoplasm and chloroplasts of mesophyll cells adjacent to the aphid stylets tracks (Fig. 2B-D, 3A-B), whereas the pre-immune sera (dilution 1:50) labelled uniformly across different compartnemtns at low level (Fig. 3C). In sections of uninfested control samples, the Mp10 antisera (1:100) did not label any compartments above background level (Fig. 3D). Thus, despite some aspecific labelling of plant proteins on immuno-blots (Fig. 1), the labelling of plant tissue adjacent to the aphid feeding site is specific. Quantification of the labelling seen on micrographs revealed consistently more labelling with Mp10 antisera of the cytosol and chloroplasts with Mp10 antisera in mesophyll cells adjacent to aphid feeding sites compared to the two control treatments (Fig. 3G, Supplementary Fig. S3). Labelling of mesophyll cells distal to the site of feeding was not detected (Fig. 3E; Supplementary Fig. S3). Results shown are representative of two independent experiments. Chloroplasts are known to be prone to high levels of non-specific background labelling in IGL experiments, however we do not see similar levels of labelling of plastids in the uninfested or preimmune controls, or of plastids distal to the site of feeding. Therefore, these data suggest that Mp10 localizes in the cytosol and chloroplasts of mesophyll cells adjacent to the stylets tracks.

**Figure 3.**
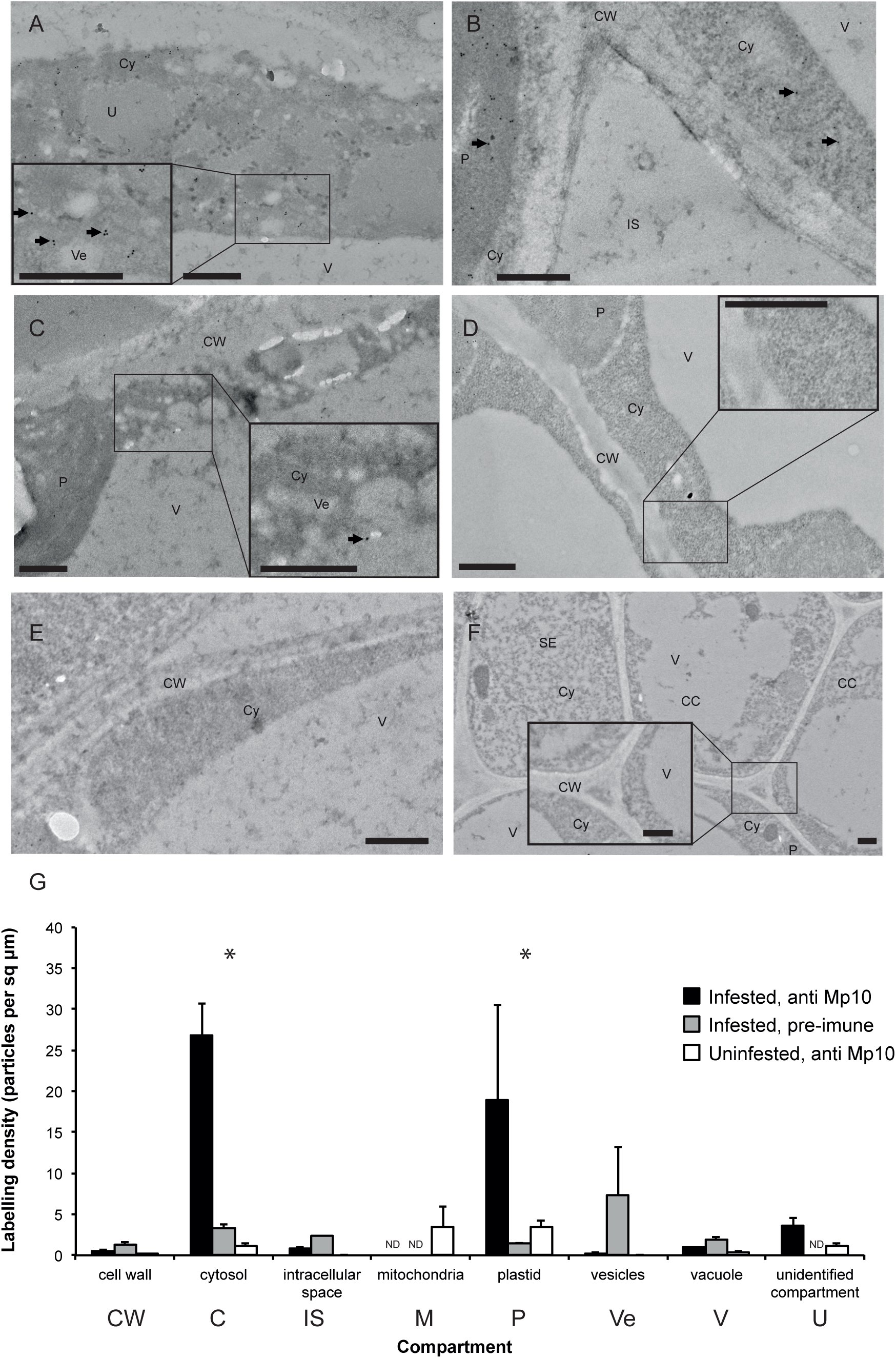
Mp10 detection is specific and restricted to the cells adjacent to stylets tracks. Ultra-thin sections from *A. thaliana* leaves infested with GPA, and labeled with anti-Mp10 (1:100 dilution) show labelling (examples indicated with arrows) in the cytosol “Cy” (A-B, G) and plastid “P” (B, G) of cells adjacent to the stylets track, but not when labelled with pre-immune serum (1:50) (C, G). Samples from uninfested plants are not labelled with anti-Mp10 above background levels (D, G). Labeling with anti-Mp10 was not detected in mesophyll cells distal to the feeding site (E), or an area of the vascular bundle closest to the feeding site in GPA-infested tissues (F). Quantification of labelling density (G) based on measurements of compartment area and gold particle count across multiple images of different areas from the samples. Values show mean (+/− SEM) density (infested, antiMp10 (at 1:100 dilution) n=13; infested, pre-immune (1:50) n=4, uninfested anti-Mp10 (1:100) n=5). Asterisks indicate significant differences in labelling both between infested vs. uninfested samples, and between anti-Mp10 and pre-immune labelled samples for that compartment (*P*<0.01, GLM). Abbreviations: “IS” intercellular space; “Cy” cytoplasm; “V” vacuole; “CW” cell wall; “Ve” vesicles; “P” plastid; “U” unidentified compartment. In (F): “CC” is a phloem companion cell, and “SE” are phloem sieve elements. Insets (A, C, D and F) are magnified regions of the image, all scale bars are 500 nm.

Occasionally we found that cells adjacent to the stylets’ track contained unusual features (Fig. 3 A, C). The cytoplasm of these cells showed reduced granularity and a higher number of vesicle-like structures that may be endoplasmic reticulum (ER)-derived. These were not seen in all aphid-infested samples (e.g. Fig. 3 B), but may represent a response of the cell to aphid probing.

Because GPA feeds from the phloem, we quantified Mp10 antisera labelling of the vascular bundles closest to the aphid feeding site (Fig. 2 A; Fig. 3F). We did not detect specific labelling of any cell type in the vascular tissue in experimental compared to the control samples (Supplementary Fig. S2B) suggesting that Mp10 is not delivered into the vasculature or is not present at high enough concentrations for detection in these tissues.

We also investigated labelling with antisera to MpOS-D1, a family member of Mp10, in and near stylets tracks. We did not detect specific labelling of mesophyll or vascular tissues with MpOS-D1 antisera in or near GPA feeding sites compared to controls (Supplemental Fig. S2C, D). While the Mp-OSD1 antisera detected denatured MpOS-D1 protein on immuno-blots (Fig. 1), it is possible that these antisera do not detect this effector in resin sections. Alternatively, GPA may not deliver Mp-OSD1 or delivers this effector at low concentrations or to different locations inside plant tissues compared to Mp10.

### Aphid effector MpPIntO1 localizes to the sheaths surrounding aphid stylets at feeding sites

To compare localizations of aphid effectors, we also conducted IGL with antisera to MpPIntO1 and MpC002 on ultra-thin sections of tissues containing aphid stylets tracks. We observed very strong and specific labelling of the stylets sheaths by the MpPIntO1 antisera (Fig. 4B; Suppl. Fig. 2G), but not by the corresponding preimmune serum (Fig. 4C; Suppl. Fig. 2G).The MpC002 antisera also labelled the sheaths more strongly than other tissues in the plant and this labelling was stronger than detected with the corresponding pre-immune sera (Fig. 4D-E; Suppl. Fig. 2E). However, the background labelling of uninfested plant tissue by the MpC002 antisera was much higher than for MpPIntO1, suggesting that anti-MpC002 may be less specific (Suppl. Fig. 2E-H). These data suggest that MpPIntO1, and possibly also MpC002, are found in the sheaths that surround the stylets during feeding.

**Figure 4.**
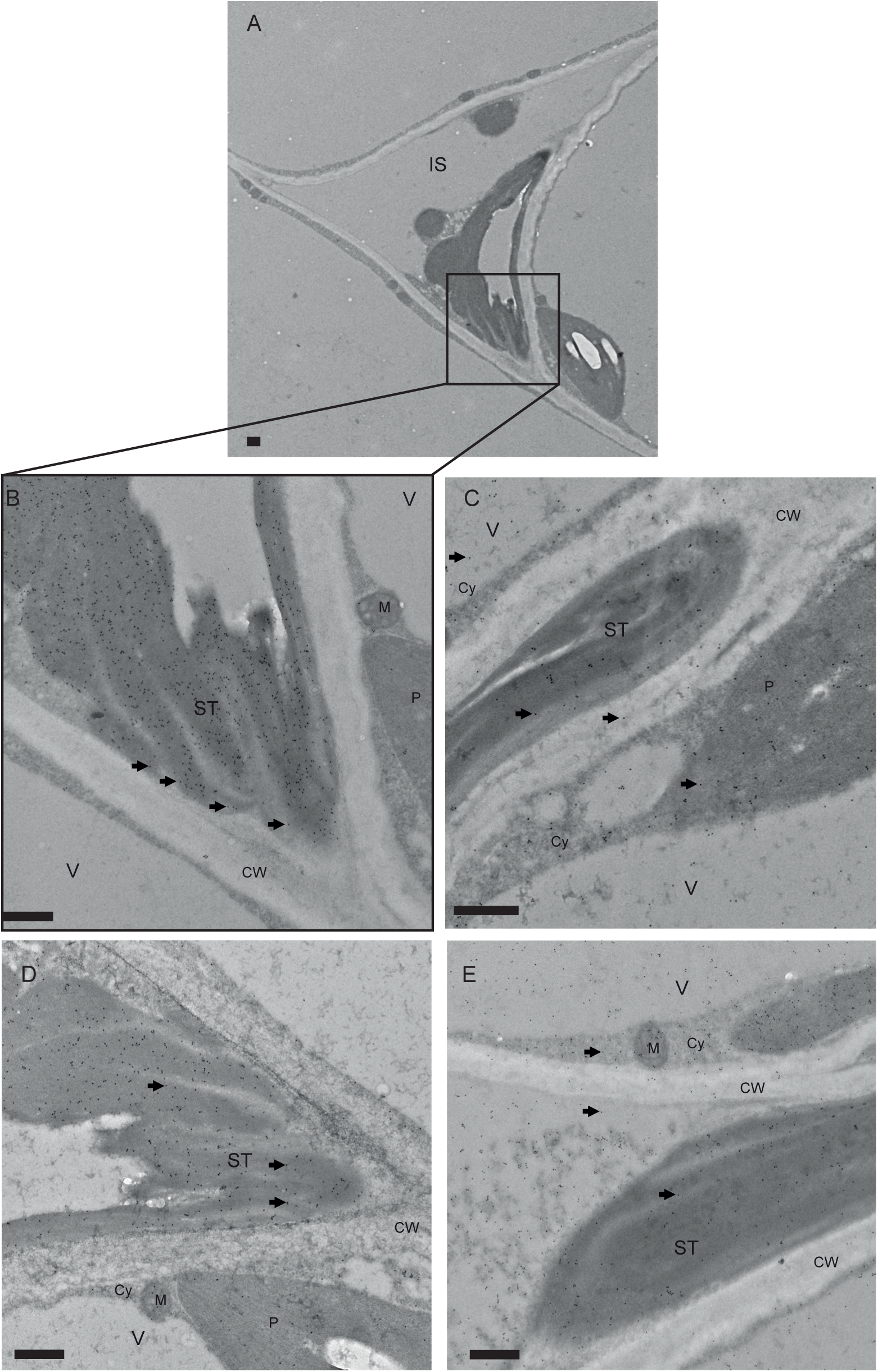
MpPIntO1 and MpC002 antisera label the stylets sheaths. Ultra-thin sections from aphid-infested samples showing the gelling saliva that forms the stylets sheath (“ST”), immuno-gold labelled with anti-MpPIntO1 (A-B at 1:100 dilution) or with the corresponding pre-immune serum (C, 1:50); or with anti-MpC002 (D, 1:100) or the corresponding pre-immune serum (E, 1:50). Examples of gold-labelling, where present, are indicated with arrows. Compartments are indicated as: “ST” stylet track; “IS” intercellular space; “Cy” cytoplasm; “V” vacuole; “CW” cell wall; “Ve” vesicles; “P” plastid; “M” mitochondria. Scale bars all indicate 500 nm. Quantification of labelling density can be found in Supplemental Fig. 3.

### Discussion

In this study, we generated specific antisera to the GPA proteins Mp10, MpOS-D1, MpC002 and MpPIntO1, which were previously shown to have activities characteristic of effector proteins (Mutti et al 2008; Bos et al. 2010; Pitino et al. 2011; Pitino and Hogenhout 2013), and used these antisera in immuno-gold labelling experiments to examine if GPA secretes effector proteins into plants during feeding. We found that the Mp10 antisera specifically labelled the cytoplasm and chloroplasts of mesophyll cells adjacent to the aphid stylet tracks. No such labelling was observed in cells further distal to the stylets tracks. In addition, immuno-gold labelling experiments with the antisera to the three other effectors did not label these compartments. Finally, we observed no or much reduced labelling densities in the pre-immune sera and uninfested controls. Taken together, these experiments provide strong evidence for Mp10 delivery by secretion in the watery saliva by GPA into mesophyll cells, where Mp10 resides in the cytosol and the chloroplasts.

Mp10 and MpOS-D1 are members of the chemosensory protein family that were first identified in the haemolymph of sensory organs. However, it is emerging that members of this family are expressed across many non-sensory tissues and may have diverse roles in processes other than chemoreception including pheromone delivery and development (Kitabayashi et al 1998, Maleszka et al 2007, Guo et al 2011), and in the case of Mp10 modulation of host plant immunity (Bos et al., 2010; Rodriguez et al. 2014). CSPs have cental hydrophobic channels that are thought to bind small ligands (Lartigue et al 2002, Mosbah et al 2003). Interestingly, the saliva of mosquitoes and other blood-feeding dipterans contain D7 proteins that are structurally related to members of the arthropod odorant binding protein (OBP) superfamily and, like CSPs, are small chemosensory proteins with globular structures and ligand-binding central hydrophobic channels (Calvo et al., 2006; Calvo et al. 2009). The D7 protein family members are anti-inflammatory mediators by binding host amines and leukotrienes resulting in, for example, inhibition of collagen-mediated platelet activation (Alvarenga et al. 2010). Thus, Mp10 may modulate plant defense responses by binding small immunosuppressive plant metabolites.

We previously found that Mp10 suppresses the flg22-induced ROS burst (Bos et al. 2010) which is dependent on BAK1 (Chinchilla et al. 2007). The ROS burst seen in response to aphid elicitors is also dependent on BAK1 (Chaudhary et al. 2014; Prince et al. 2014). BAK1 is a key regulator of several plasma membrane-localized pattern recognition receptors (PRRs), which sense potential invaders early on and trigger downstream immune signalling. Thus, given that we found Mp10 in the cytoplasm of mesophyll cells adjacent to stylets tracks, this effector may suppress the ROS burst induced by aphids shortly after initial probing by the aphid in the pathway phase. The additional detection of Mp10 in chloroplasts is of interest because chloroplasts are a major sources of ROS and have a critical role in PTI (Shapiguzov et al. 2012; Nomura et al. 2012; Caplan et al. 2015). Given that Mp10 does not appear to have an obvious plastid transit peptide sequence, it is unclear how this protein gets into chloroplasts. As a small protein, Mp10 may migrate into chloroplast passively or via interaction with ligands that are located predominantly in these cell organelles.

In addition to the suppression of PTI, heterologous expression of Mp10 in plants elicits an effector-triggered immunity (ETI)-type response, leading to chlorosis and activation of JA- and SA- signalling pathways (Bos et al. 2010; Rodriguez et al. 2014). However, in these expression experiments Mp10 is present at high levels in many cells, whereas our data indicate that Mp10 is delivered in only some cells near the aphid stylets in the presence of other effectors (such as MpPIntO1). Hence, it remains to be investigated how Mp10 modulates cell defense responses during aphid feeding.

Our IGL data show different localizations of MpPIntO1 and MpC002 compared to Mp10. MpPIntO1 and MpC002 are abundant proteins in aphid saliva and can be readily detected by protein immuno-blotting in artificial diets fed upon by aphids and extracts of aphid-exposed leaves (Mutti et al. 2008; Harmel et al. 2008). Our IGL data provide evidence that MpPIntO1, and possibly also MpC002, are present in the stylets sheaths. We did not detect MpPIntO1 inside plant mesophyll and vascular cells indicating that aphids do not deliver this effector inside these cells or else they are present at or below detection-level concentrations. Our data are in agreement with previous findings showing that MpPIntO1 is abundant in the gelling saliva, which generate the sheaths (Harmel et al. 2008), though MpPIntO1 and MpC002 are also found in soluble saliva in artificial diets fed upon by aphids (Harmel et al., 2008; Will et al., 2012; Chaudhary et al. 2015). However, given that both MpPIntO1 and MpC002 are abundant, it is possible that some of these proteins are not captured during the gelling process and end up in the soluble saliva fraction. We previously found that aphids reproduce better on transgenic *A. thaliana* that produce MpPIntO1 and MpC002 under control of 35S promoters (Pitino and Hogenhout 2013), though the effect of MpPIntO1 on aphid performance is variable (Pitino and Hogenhout 2013; Elzinga et al. 2014). In addition, RNA interference (RNAi)-mediated knock down of MpC002 in aphids reduced aphid performance on plants, whereas MpPIntO1-RNAi aphids were not affected (Pitino et al. 2011). Several pathogens deliver effectors in the apoplastic space to modulate defence responses (Tian et al. 2004, 2007; De Wit 2016). Our IGL data open up the possibility that the aphid MpPIntO1 and MpC002 effectors act in the plant apoplast from within sheath saliva.

In some cases, changes in subcellular structures occurs in response to aphid feeding (Fig. 3, A, C). It is known that pathogen attack can induce dramatic reorganisations of subcellular structures, including organelles (Hardham et al. 2008; Ben Khaled et al. 2015). Aphids can also cause significant tissue damage and disruption of plant tissue during feeding (Saheed et al. 2007). Moreover, probing of cells by aphid stylets induces a rapid subcellular relocalization of vesicle-associated *Cauliflower mosaic virus* (CaMV) particles that is essential for the acquisition and transmission of this virus by aphids (Martiniere et al. 2013). The increase in abundance of vesicular structures we observed might be associated with the activation of plant defence responses, such as the delivery of defence compounds via vesicles to deter attack. Alternatively, aphid effectors may modulate plant cell processes, such as the reprogramming of vesicle trafficking, as has been shown to occur for pathogens (Ben Khaled et al. 2015; Bozkurt et al. 2015).

## Materials and methods

### Antibody generation

Coding sequences corresponding to the predicted mature effector proteins (minus predicted secretory signal peptide) were expressed in *Escherichia coli* as *N*-terminal 6Xhis tag fusions (Mp10, MpPIntO1 and MpOSD1) or as a *N*-terminal dC, and 6XHis tag fusion (MpC002), purified and checked for purity by protein immuno-blotting using anti-His-tag antibody (BacPower and FoldArt technologies, Genscript). Proteins were injected into chicken (Mp10) or rabbit (MpOSD1, MpPIntO1, and MpC002) by Genscript. Specific antisera were affinity purified using immobilized recombinant protein. Aliquots of pre-immune serum were collected from the animals before injection.

### Western blotting

*Arabidopsis thaliana* Col-0 plants were exposed to GPA (*Myzus persicae* clone O) for 24h (1000 aphids per 3-week old plant), aphids were carefully removed, and the plant rosette was rinsed in distilled water to remove any remaining aphid material from the surface. Total protein was extracted from plant and aphids in sodium dodecyl sulfate (SDS) buffer (5 µl per mg plant tissue, or 200 µl per mg aphid tissue), and 10 µl was size-separated by SDS-polyacrylamide gel electrophoresis and blotted onto nitrocellulose membrane using standard methods. Blots were probed overnight with 1:100 or 1:1000 antibody dilutions (or with pre-immune sera at double the concentrations), and a secondary antibody-horseradish peroxidase-conjugate.

### Immuno-localisation

Samples were harvested from aphid-infested (or uninfested) *A. thaliana* plants and immediately fixed in 4% formaldehyde / 0.5% glutaraldehyde in phosphate-buffered saline (PBS) at 4 degrees overnight. The samples were then embedded in LR White resin (Agar Scientific) using the progressive lowering of temperature (PLT) method using the Leica EM AFS2 (Automatic Freeze Substitution) machine (Caillaud et al. 2014). Sections were prepared on a Leica UC6 ultramicrotome. Sections of 0.5 µm were stained with toluidine blue for light microscopy (Nikon Microphot-SA) and ultrathin sections of approximately 90 nm, were picked up on to pyroxylin- and carbon-coated gold grids (Agar Scientific) for immuno-labelling.

The ultrathin sections were immuno-gold labelled according to the following protocol: 50mM glycine/PBS for 15 mins; Aurion blocking buffer (Aurion, The Netherlands for 30 mins; 0.1% acetylated bovine serum albumin (BSA-C) (Aurion, The Netherlands) in PBS for 2 × 5 mins; primary antibody of choice at 1:10 to 1:100 dilutions in 0.1% BSA-C/PBS for 90 mins; 0.1% BSA-C/PBS for 6 × 5 mins; secondary antibody conjugated to 10nm gold particles (1/40 dilution in 0.1% BSAC/PBS) for 90 mins; 0.1% BSA-C/PBS for 6 × 5 mins; PBS for 2 × 5 mins; and water for 2 × 3 mins.

Grids were viewed in a FEI Tecnai 20 transmission electron microscope (FEI UK Ltd, Cambridge, UK) at 200 kV and imaged using an AMT XR60 digital camera (Deben, Bury St Edmunds, UK) to record TIF files. Images were analysed using ImageJ. The areas and the numbers of gold particles detected in each subcellular compartment were measured across a series of images across each sample. The density across different compartments in infested/uninfested samples were analysed by general linear model (GLM, Genstat). Where differences in labelling intensity were detected between aphid-infested and uninfested samples, replicate sections were probed with the corresponding pre-immune serum, at a higher concentration that the antibody was used at, to demonstrate that labelling was specific.

## Acknowledgements

We thanks the members of the Hogenhout lab for useful discussions and feedback, and the BBSRC for funding (BB/J0045531/1).

---

**Supplementary Figure 1.**
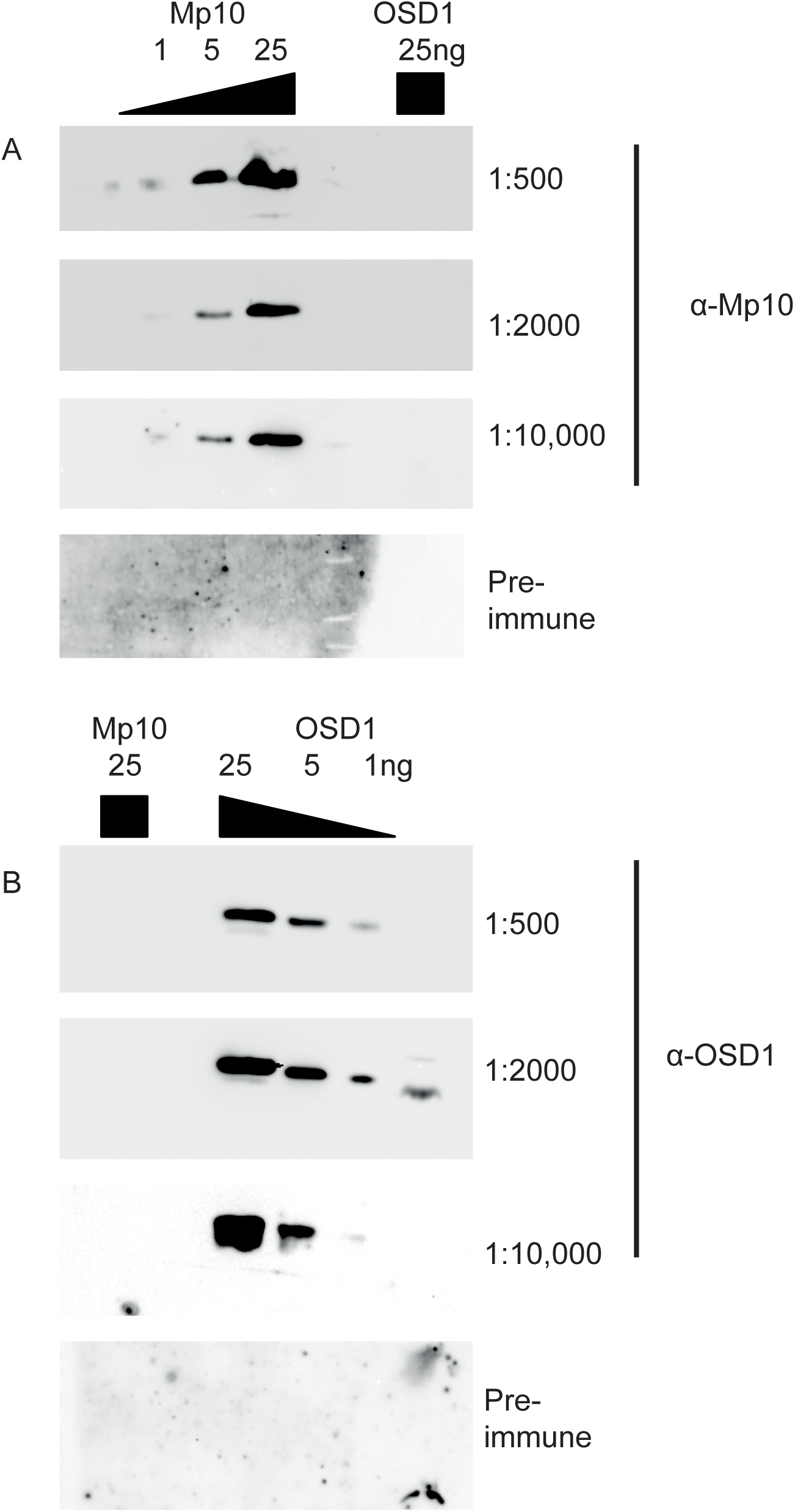
Mp10 and MpOSD1 antisera are specific and do not cross react. Immuno-blots of pure recombinant Mp1\0 and MpOSD1 proteins probed with antisera raised against the proteins show the sensitivity and specificity of these antisera. (A) Replicate blots of a range of quantity of Mp10 protein (1-25ng) and 25ng OSD1 protein probed with (from top) decreasing concentration of the anti-Mp10 serum (from 1:500 to 1:10,000) or with corresponding pre-immune serum (at 1:1000 dilution). (B) Replicate blots of a range of quantities of OSD1 protein (1-25ng) and 25ng Mp10 protein probed with (from top) decreasing concentration of the anti-OSD1 serum (from 1:500 to 1:10,000) or with corresponding pre-immune serum (at 1:1000 dilution).

**Supplementary Figure 2.**
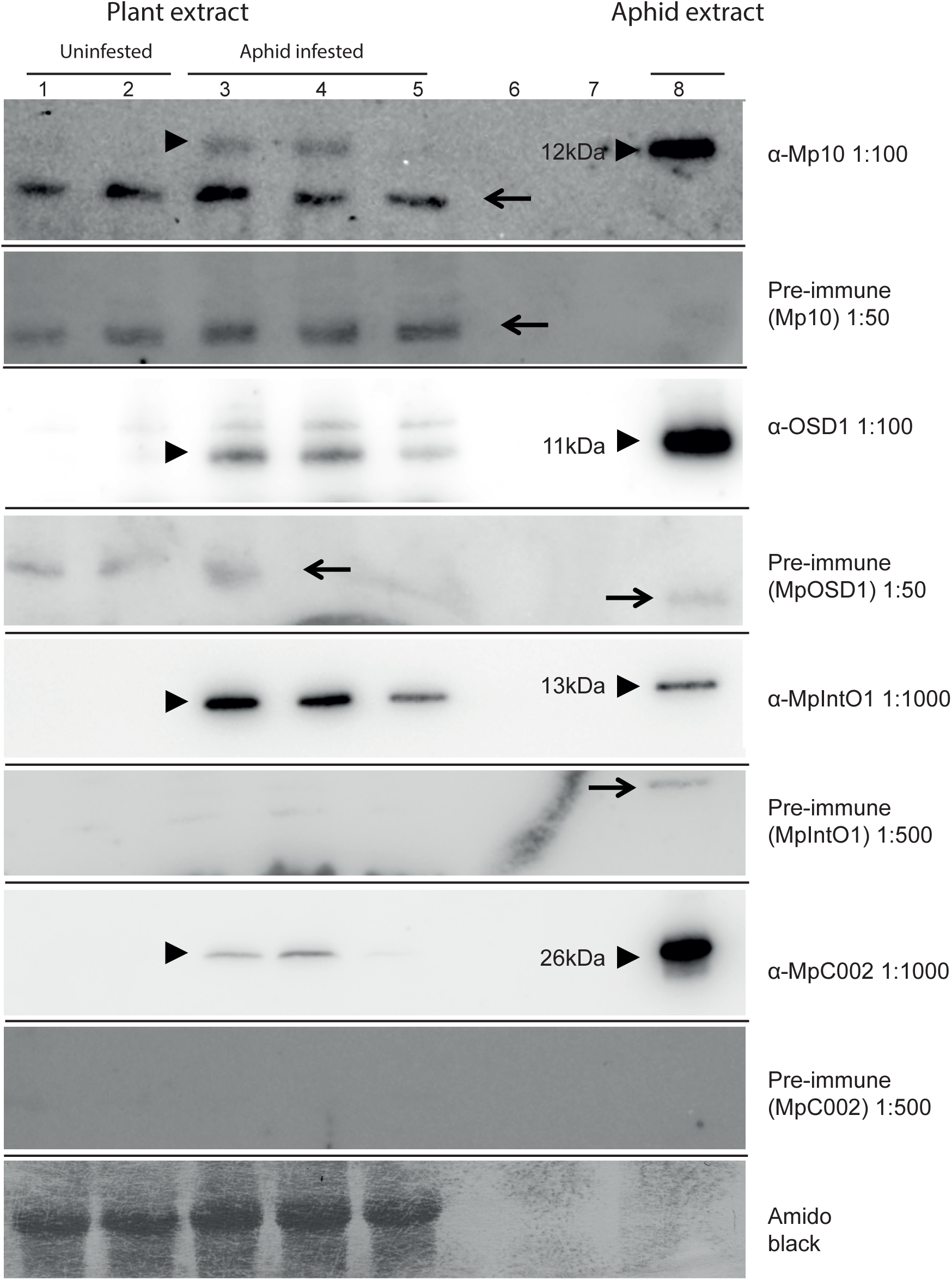
Immuno-blots of aphid-exposed plant tissue probed with antisera raised against candidate effector proteins and also with corresponding pre-immune serum. Immuno-blots of total protein extracts from Arabidopsis rosettes; uninfested (lanes 1-2) or infested with GPA (lanes 3-5), and extract from GPA aphids (lane 9), were probed with antisera raised against the aphid salivary proteins (as shown in Fig. 1), and with the corresponding pre-immune sera (PI); (from the top) Mp10, Mp10-PI, OSD1, OSD1-PI, MpPIntO1, MpPIntO1-PI, MpC002, and MpC002-PI. The bottom panel shows loading of the plant extract samples by amido-black staining of the membrane. Arrowheads indicate bands of the expected sizes detected in aphid-exposed plants and aphids, but not unexposed plants. Arrows indicate presumably non-specific bands of other sizes. Pre-immune sera were used at higher concentrations than all corresponding antibodies to ensure that specific antibody-labelled bands were not also detected by pre-immune serum.

**Supplementary Figure 3.**
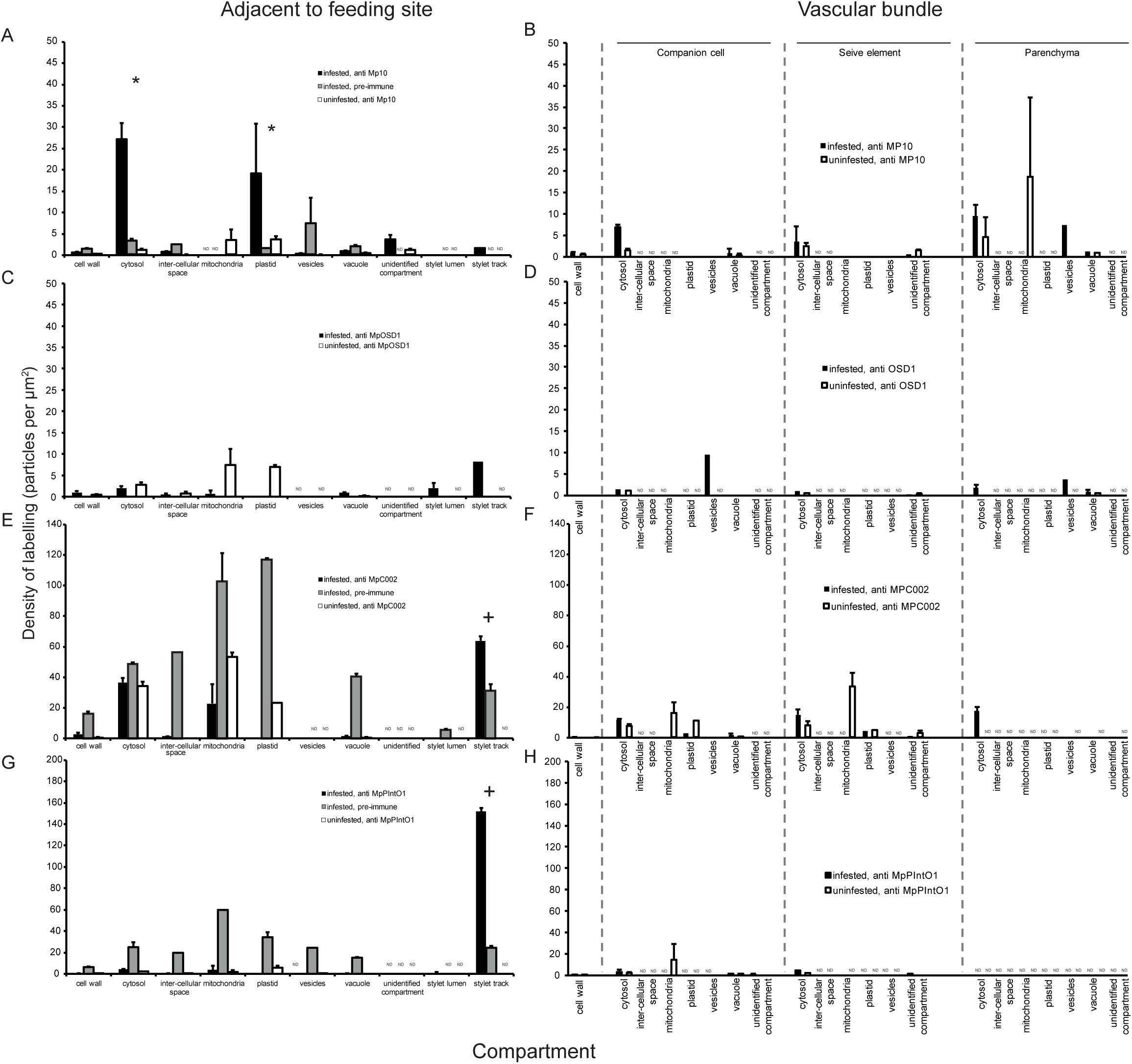
Quantification of immuno-gold labelling over different cellular compartments and tissues. Graphs show the density of gold particles in different subcellular compartments (mean +/− SEM (n= from 3 to 13)) for (left) cells adjacent to the aphid stylet track (or equivalent mesophyll cells from uninfested tissues), and (right) cells in the vascular bundle. Samples were probed with affinity-purified antibody (all at 1:100 dilution) raised against Mp10 (A-B); MpOSD1 (C-D); MpC002 (E-F) or MpPIntO1 (G-H), or (where indicated) with the corresponding preimmune serum (all at 1:50 dilution). Asterisks indicate significant difference in labelling both between infested and uninfested samples, and between antibody and pre-immune labelled samples for that compartment (*p* < 0.01, GLM). Plus signs indicate significantly higher labelling of stylet track relative to other compartments in infested tissue (for which no uninfested control plants could be tested) and for which the signal was significantly higher than for infested samples probed with the corresponding pre-immune serum (*p* < 0.01, GLM). “ND”: Not determined.

